# Spatial heterogeneity in collective electrotaxis: continuum modelling and applications to optimal control

**DOI:** 10.1101/2024.02.28.580259

**Authors:** Simon F. Martina-Perez, Isaac B. Breinyn, Daniel J. Cohen, Ruth E. Baker

## Abstract

Collective electrotaxis is a phenomenon that occurs when a cellular collective, for example an epithelial monolayer, is subjected to an electric field. Biologically, it is well known that the velocity of migration during the collective electrotaxis of large epithelia exhibits significant spatial heterogeneity. In this work, we demonstrate that the heterogeneity of velocities in the electrotaxing epithelium can be accounted for by a continuum model of cue competition in different tissue regions. Having established a working model of competing migratory cues in the migrating epithelium, we develop and validate a reaction-convection-diffusion model that describes the movement of an epithelial monolayer as it undergoes electrotaxis. We use the model to predict how tissue size and geometry affect the collective migration of MDCK monolayers, and to propose several ways in which electric fields can be designed such that they give rise to a desired spatial pattern of collective migration. We conclude with two examples that demonstrate practical applications of the method in designing bespoke stimulation protocols.

## 1 Introduction

During electrotaxis, the process by which eukaryotic cells establish a cell polarity and move directionally in the presence of an electric field [1–3], the electric field acts as an external driver of collective cell migration and has a dramatic impact on collective behaviour by providing the constituent cells in a tissue with a persistent cue to migrate in a given direction. For this reason, electrotaxis has become known as a method to reliably ‘steer’ a cellular collective [3]. In previous work, we explored the possibilities for optimal steering of an epithelial monolayer [4] by considering the velocity of the centre of mass of the monolayer, and deriving an optimal stimulation protocol such that the velocity of the centre of mass satisfies certain optimality conditions. This work seeks to refine that approach by considering the question of how to use electric fields to steer entire tissues. That is, rather than controlling just the movement of the centre of mass, how can mathematical modelling be used to design electric fields to move entire tissues, making sure that the entire tissue moves according to some desired velocity?

Controlling the movement of an entire cellular collective is significantly more challenging than controlling the movement of its centre of mass, for the following reason. In a large epithelial monolayer, there exist several endogenous cues that lead cells in different positions to have a preferential polarity, and exert active forces in different directions. For example, in MDCK monolayers there exists a clear polarity at the edges that acts as a cue for cells to migrate outward [1,5,6]. Therefore, when studying how to use an electric field to steer an entire epithelial monolayer, one must have a good understanding of how the endogenous migratory cues inside the monolayer compete with the external migratory cue of the electric field to determine the cell polarity and active forces that drive collective migration.

The question of how the electric field interferes with local endogenous cues has been documented experimentally but is still not well understood. For example, Wolf *et al*. find that the electric field reprograms long-term supracellular dynamics, and that the edge regions of the tissue have significantly different velocity dynamics than the bulk of the tissue [7]. Modest theoretical advances exist to understanding mechanistically how the electric field interferes with endogenous migratory cues to create collective migration. For example, Zhang *et al*. [8] develop an on-lattice model that phenomenologically models the collective response to the electric field as a linear superposition of different forces, including cell-cell repulsion, and a force in the direction of the electric field. While these results can capture some of the features of collective electrotaxis, they do not take into account the temporal dynamics of adaptation to the field, which, as we demonstrated in [4], have a dramatic effect on the controllability of the tissue. Likewise, the on-lattice model cannot be used directly within a formal control framework to design optimal electric fields for any given velocity pattern, which makes the model difficult for practical applications in tissue engineering. To overcome both these difficulties, this work will be concerned with developing robust continuum models that can capture the spatial heterogeneity of velocities in MDCK monolayers undergoing electrotaxis, and using these models to propose the design of electric fields that induce collective migration.

As a first contribution, this work will propose an extension to the ordinary differential equation (ODE) model for the tissue bulk velocity proposed in [4]. This ODE model, inspired by an *adaptation-excitation model* proposed by Erban and Othmer [9], models how cells are *excited* when initially exposed to an external signal, and how they slowly adapt when exposed for long periods of time. The spatial model extension considered in this work will introduce terms that relate the position of cells in the monolayer to endogenous cues and their sensitivity to the field. This model extension is needed because experiments by Cohen *et al*. [1] suggest that cells at the tissue edges appear to be less responsive to the field than cells in the bulk. How this weaker sensitivity to the field then interacts with the endogenous cues inducing outward migration at the tissue edges is not understood. Therefore, to understand the spatial and temporal heterogeneity in the responses of the tissue to electric field stimulation, there is a need to have detailed and reliable modelling of how cells integrate cues. Our model extension shows that the the migratory cues at the tissue edges can be represented as the superposition of a fixed cue pointing out of the tissue, and an electrotactic cue that is weaker than the cue in the tissue bulk.

Still, this model extension uses a system of coupled ODEs to model competing migratory and electrotactic cues in distinct, but fixed, tissue regions. However, for bioengineering applications we are generally interested in predicting the velocity during electrotaxis throughout the entire monolayer, regardless of tissue shape. Therefore, we take the insights from this ODE continuum model of cue competition in different tissue regions, and develop a novel reaction-advection-diffusion model for collective electrotaxis that can predict the movement of an entire MDCK monolayer as it undergoes electrotaxis. This continuum model can readily be used to design bespoke electric fields for desired migratory patterns. We also use the model to predict how tissue size and geometry affect the collective migration of MDCK monolayers during electrotaxis.

This work is structured as follows. In Section 2 we briefly present the data from Wolf *et al*. [7] on the electrotactic response in different regions of MDCK epithelial monolayers and introduce some of the most notable findings regarding the experimentally observed spatial heterogeneities. We proceed in Section 3 by analysing the question of how different migratory cues are interpreted together to give rise to different velocity distributions in different regions in the tissue, using a system of coupled ODEs to model competing migratory and electrotactic cues in distinct, but fixed, tissue regions. In order to model the velocity of electrotaxis throughout the entire monolayer, regardless of tissue shape, we use the insights gained from our ODE model in Section 3 to derive and validate a diffusion-advection-reaction equation that describes the movement of an epithelial monolayer undergoing collective electrotaxis. We use this continuum model to propose several ways in which electric fields can be designed such that they give rise to a desired spatial and temporal pattern of collective migration.

## 2 Experimental data and methods

Experimental data are publicly available from Wolf *et al*. [7]. In these experiments, a total of nine MDCK epithelial monolayers were grown to confluence and stimulated with an electric field of 3V/cm. Velocities during the experiment were computed using particle image velocimetry (PIV). PIV data was acquired in each of the different tissue regions in Figure 1 (left), and PIV data from the experiment for each tissue are shown in Figure 1 (right). We remark that only the top edge is shown for the two edges parallel to the electric field direction, since their velocity dynamics are identical [7]. For the computation of local cell densities, we used the segmented cell nuclei data available from Wolf *et al*. [7] and computed two-dimensional histograms with a grid of size 150*×*150. A one-dimensional density profile, averaged in the *y*-direction, in the leading and trailing edge of a representative tissue, is shown in Figure 1 (bottom row).

**Figure 1:**
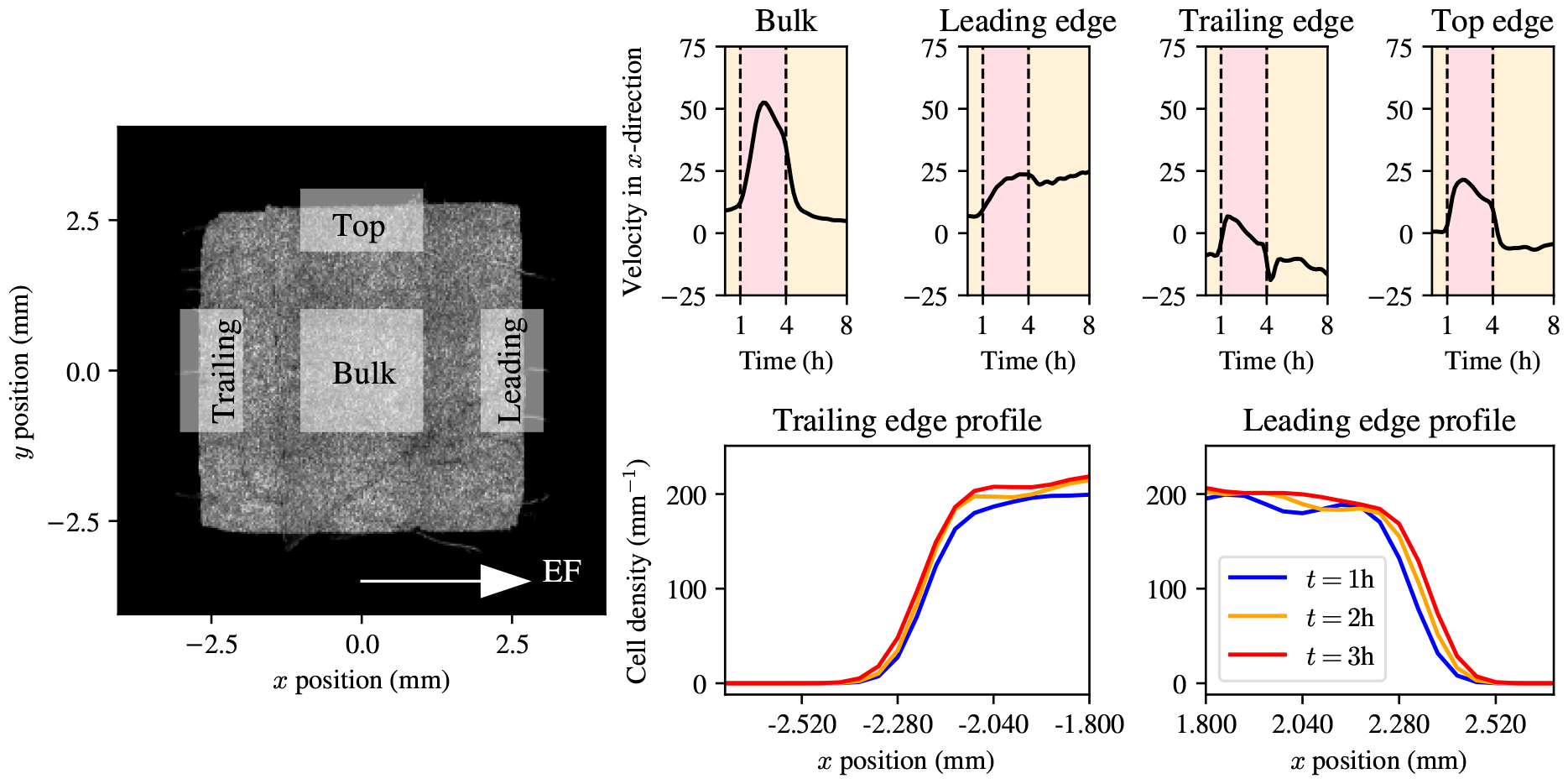
Collective electrotaxis data from Wolf *et al*. [7] panel: phase image of a representative square MDCK-II epithelial monolayer, with different tissue regions marked. Top row: average velocity data in the bulk, leading edge, trailing edge, and top edge, respectively. Red shaded times indicate electric field on. Bottom row: cell density profile in space at three time points during the experiment.

The experimental velocity data in Figure 1 (right) contain a number of notable features. The first notable feature is that each of the edges has its own representative behaviour, with the trailing, leading, and top regions each having a different velocity trace, which are all distinct from the bulk. The second notable feature is that the edges of the tissue all exhibit a far smaller velocity during electrotaxis than the tissue bulk. Interestingly, all the edge regions experience a slowdown during electrotaxis over a time scale similar to that of the slowdown in the bulk. The final notable feature is that both the top edge and the trailing edge seem to experience *recoil* after the electric field is turned off. That is, for a period of time after the electric field is turned off, the velocity in the top edge is negative, and the velocity in the trailing edge exhibits a sharp undershoot with respect to its velocity during electrotaxis and free epithelial expansion. This finding was also reported by Wolf *et al*. [7], who related it to mechanical factors. In the remainder of this work, we will explore these three phenomena and investigate the possible mechanisms responsible for creating them.

## 3 Modelling spatial heterogeneity during electrotaxis

Having understood that the edge regions of the tissue exhibit velocity dynamics that are different from those of the bulk of the tissue, in this section we propose an extension to our previous work which proposed an ODE model for the bulk velocity of an electrotaxing epithelium [4] to account for the observed heterogeneity in velocities between the different regions of the tissue.

In Section 3.1 we propose the model extension as a system of ODEs describing the response in the different tissue regions. In Section 3.2 we perform Bayesian inference on the model, with a view to understanding how well the model can describe the experimental data.

### 3.1 An ODE model for vector superposition of competing migratory cues

Figure 1 shows that different regions of the epithelial monolayer migrate at different speeds during electrotaxis: the bulk of the tissue migrates at the highest speed of all tissue regions; and the trailing and top edges migrate slower than the leading edge. The fact that different regions of the monolayer migrate at different speeds should not be surprising, given that each tissue region is subject to different migratory cues: MDCK-II epithelial monolayers are well-known to exhibit persistent spreading behaviours, even in the presence of electrotactic or chemotactic cues [1,3,7]. As such, there is competition between different cues inducing migration in different directions at every tissue location, and tissue migration is the result of how these cues are interpreted together.

In order to construct a continuum model of collective migration during electrotaxis, we first seek to understand the mechanisms driving these differences in observed migratory velocity. We do this by modelling the average velocity of the different tissue regions using an extension of the one-dimensional ODE model for the velocity in the direction of the electric field developed, calibrated, and optimised in our earlier work [4]. In brief, the model considered in [4] and based on Erban and Othmer [9] considered the electric field, *s*, as an external stimulus. It describes the cellular response by modelling two chemical species that dictate intracellular signalling: an *effective signal, s*_eff_, and an *inhibitor, I*. The effective signal, *s*_eff_, is a scalar quantity that describes the extent to which an external signal is transmitted within the cell [4]. There is therefore a difference between the constant applied electric field applied and the internal signal. The inhibitor, in contrast, represents the strength of the internal signalling pathways that try to inhibit this response to the external electric field stimulus. The dynamics are given by

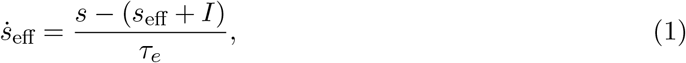

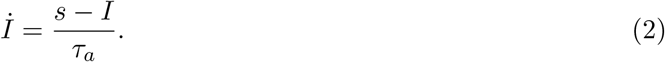

In this model, one assumes that the excitation timescale, *τ*_*e*_, is much shorter than the adaptation timescale, *τ*_*a*_. Finally, assuming that whenever *s ≥* 0, the effective signal, *s*_eff_, equally satisfies *s*_eff_ *≥* 0, we consider the quantity

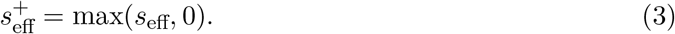

For more details on the model and its numerical implementation, we refer the reader to [4]. To extend the model in Equations (1) and (2), we formulate a phenomenological ODE model for the velocity in the direction of the electric field in each of the edge regions. Previously, the velocity of the bulk of the tissue, *v*, was modelled by describing the average acceleration of the bulk of the tissue as a balance of active forces in the direction of the electric field against forces resulting from friction with the substrate. In [4], we assumed that cells experience a viscous friction force per unit mass which is proportional to their velocity with a friction coefficient, *γ*, such that the friction force per unit mass is given by −*γv*. Furthermore, we assumed that the active force per unit mass in the direction of the electric field is proportional to the effective signal, 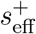 [4]. This assumption results in the following expression for the active force per unit mass, 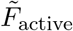,

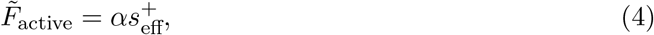

where *α >* 0 is a parameter that controls the responsiveness of the bulk to the input signal. By then considering a balance of forces per unit mass, one obtains a single equation for the velocity

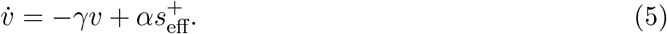

In the tissue bulk, this simple force balance is an appropriate model of the constituent forces since, on average at the continuum length scale, cells do not exert forces in any preferential direction. At the edges of MDCK-II epithelial monolayers, however, it is well-characterised that cells in a region within ∼100*μ*m of the tissue edge exert active forces in the direction of the normal at the tissue boundary [6]. For this reason, a first extension in this model of acceleration in different tissue regions is to include an active force per unit mass, **F**_edge_, in the direction of the normal at the tissue edge, *i*.*e*.,

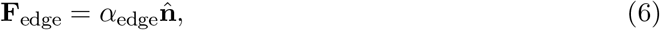

where 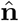 is a unit vector and *α*_edge_ is a constant that controls the magnitude of the force per unit mass. Note here that in the specific example of the square tissues considered in this study, the direction of the outward normal is constant throughout each of the edge regions.

A second extension we formulate takes into account the location of cells within the tissue, and its resulting effect on the migratory velocity. Both *in vivo* and *in vitro* cell location within a tissue plays an important role in collective migration, since the physical cues received by cells in collectives has an effect on the magnitude and the direction of active force generation [10–13]. At the tissue edges with a normal vector perpendicular to the direction of the electric field, *i*.*e*., the top and bottom edges, cells have a preferred polarity in the direction of the normal due to free-space sensing of cells at the edge, as well as contact inhibition of locomotion [1, 7, 14]. Given that cell polarity in the edge regions whose normal is perpendicular to the electric field might influence the orientation of the cytoskeleton, and this might have a direct effect on the magnitude of the active force in the direction of the electric field, we propose a model extension to take this into account. In the original model in Equation (5), the magnitude of the active force per unit mass is given by a single constant, *α*. As a model extension in this paper, we allow the magnitude of the active force per unit mass to vary, depending on the position in the monolayer. To do so, we introduce a constant for the magnitude of the field response in regions of the tissue whose cell polarity is parallel to that of the electric field, which we denote by *α*_∥_, and another for the tissue regions whose polarity is perpendicular to that of the electric field, which we denote by *α*_⊥_. Since cells in the tissue bulk have no preferential polarity at the continuum length scale, and polarise quasi-instantaneously in the direction of the electric field when it is applied [1, 7], we model active forces per unit mass in the direction of the electric field as

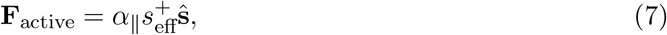

for the bulk of the tissue, where here ŝ is a unit vector pointing in the direction of the electric field, which we assume to be constant, uniaxial and spatially uniform. This framework can be naturally extended to time-varying and spatially heterogeneous electric fields. Finally, to model the apparent phenomenon that cells at the tissue edges are less sensitive to the electric field than cells in the bulk, we model that the active forces in the leading and trailing edge regions are given by

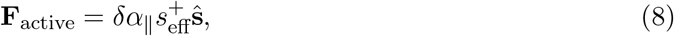

and by

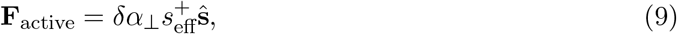

for the top region. Here, *δ* ∈ [0, 1] is a constant which scales the active force per unit mass in the direction of the electric field to take into account that cells at the tissue edges are less sensitive to the electric field. Putting the forces per unit mass in dynamic equilibrium finally yields

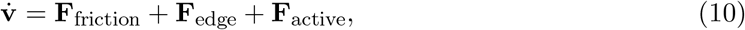

for each of the tissue regions as described above. In this case, we have assumed that the different migratory cues act additively on the monolayer regions, as we take the resultant force to be the sum of each of the active forces. As in the previous paper, we take the friction force per unit mass to be linear in the velocity. Taking components in the direction of the electric field, *i*.*e*., in the *x*-direction, gives a system of one-dimensional ODEs for the velocities in each of the different tissue regions:

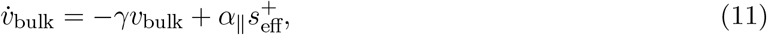

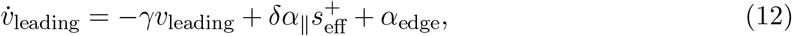

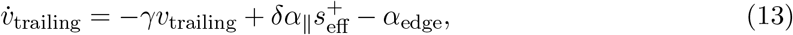

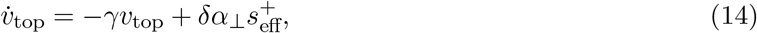

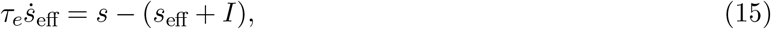

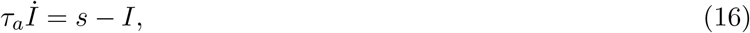

where, like in the previous paper,

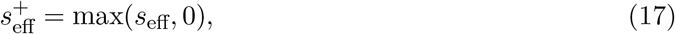

and *s* is the field strength in the *x*-direction. We reiterate that this model only includes the *x*-velocity dynamics for the top edge, since the bottom edge dynamics are identical by symmetry.

### 3.2 Bayesian inference of model parameters in the superposition model

In this section, we aim to validate the hypotheses presented in the previous section by calibrating the model in Equations (11)-(16) to the experimental data from Wolf *et al*. [7] presented in Section 2. Following [4], we note that the velocity decay parameter, *γ*, can be estimated directly from the data, since Equation (5) predicts that given an initial velocity, *v*_0_, the velocity in the bulk of the tissue decays exponentially in the absence of an electric field. Therefore, one can fit an exponential decay to the velocity in the bulk region post-stimulation (see SI Section S1). This gives an estimate of *γ* = 1.765h^−1^. Since the bulk stimulation temporal dynamics in the simple adaptation-excitation model of [4] gave predictions that were in excellent agreement with the experimental data, we equally fix the values for *τ*_*a*_ = 2.04h and *τ*_*e*_ = 0.26h to reduce the dimensionality of the parameter space. We perform Bayesian inference of the remaining parameters, *α*_∥_, *α*_⊥_, *δ*, and *α*_edge_. We use Markov chain Monte Carlo with a Haario-Bardenet adaptive covariance and four chains under the assumption of a Gaussian error implemented in Python with the PINTS pacakge [15]. As a metric of convergence we use the 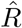 statistic [15], which summarises mixing and stationarity of the chains. We use the reference value of 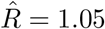 for four chains [16] and set a maximum number of MCMC iterations to 2 *·* 10^4^. The resulting posterior means are given in given in Table 1.

**Table 1:**
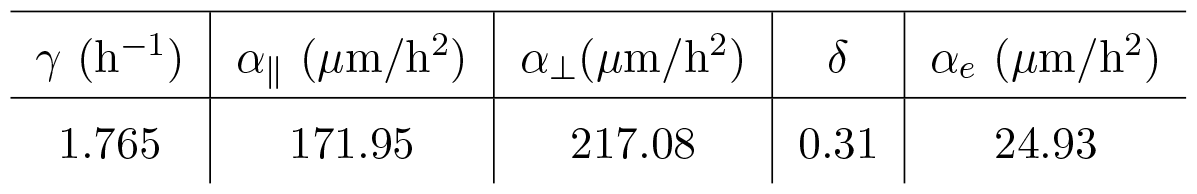
Posterior means for the extended edge velocities model, Equations (11)-(16). Note that the value for the velocity decay rate, *γ*, is estimated directly from the data.

The top row in Figure 2 shows the resulting marginal posterior distributions for these estimated model parameters. It can be seen that the marginal posterior distributions are all well-identified and concentrated around their means, showing that, given the available experimental data, the model parameters can be accurately estimated. The posterior means reported in Table 1 suggest that the three different edge regions are indeed less sensitive to the field, given the small posterior mean value of *δ*. Note also that *α*_⊥_, the strength of the active forces in the direction of the electric field, is greater than *α*_∥_, suggesting that cells with a preferred polarity that is not in the direction of the electric field have a stronger response to the electric field than the cells whose preferred polarity, and therefore also that of their competing migratory cues, is in the direction of the electric field.

**Figure 2:**
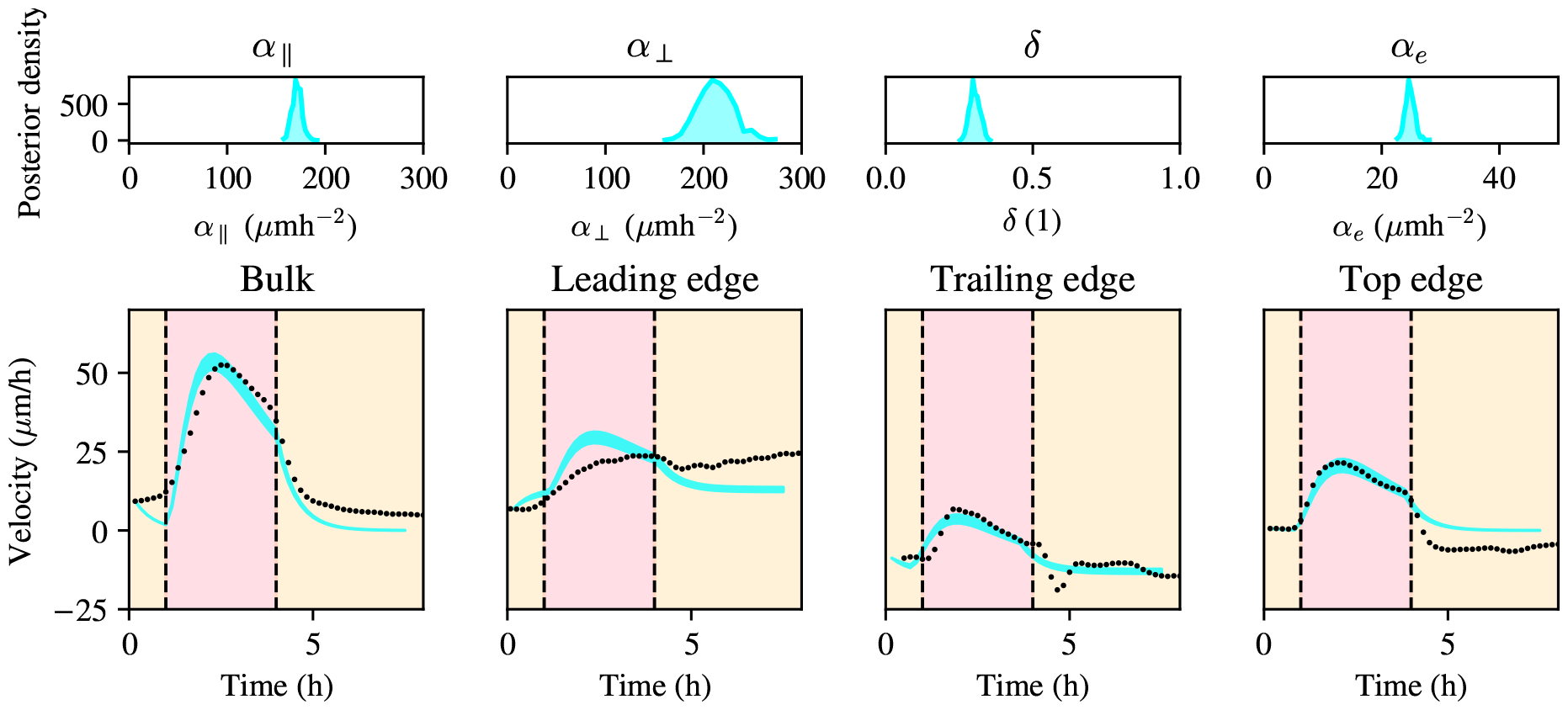
Bayesian inference with a model using vector superposition of the electric field with endogenous cues according to Equations (11)-(16). Top row: posterior distributions for model parameters *α, δ*, and *α*_*e*_, respectively, from left to right. Bottom row: posterior 95% confidence interval for the vector superposition model in cyan with experimental data (black scattered points).

To understand the validity of the model in reproducing the experimental data of Wolf *et al*. [7], and hence the underlying assumption that the edges have a weaker response to the electric field on the one hand, and that the electric field cues combine with edge endogenous migratory cues in an additive way, on the other, we calculate the posterior predictive intervals for the data resulting from the posterior distributions of the model parameters. These intervals include the middle 95% percentiles of model values found by simulating using parameters distributed according to the posterior distribution. Inspection of these posterior predictive intervals show that the model predictions are generally in good agreement with the experimental data and that the model can explain the different behaviours of the bulk of the tissue and the edges during stimulation. An important feature of the experimental data in Figure 2 is that the top edge and the trailing edge display an ‘undershoot’ of the velocity [7] after stimulation is halted, *i*.*e*. the average velocity in the direction of the field in the top edge is negative for several hours post-simulation. We comment on this phenomenon in the discussion, Section 6, where we show that it has important implications for how to model cellular transduction of the electric field signal as a function of position inside the monolayer. Finally, we remark that, at the leading edge, Figure 2 shows that the model moderately overestimates the velocity as a result of vector superposition of the electric field cue together with the endogenous cue to migrate rightward and out of the tissue. This shows us that, while the model extension presented in this section can capture large scale features, there are some finer details that are not well explained. Addressing these details will require a finer quantification of velocity, force and polarity at the monolayer edge and should be the focus of new research into collective migration during electrotaxis.

In sum, we have derived an ODE model that can describe the main features of the velocity dynamics in different, but fixed, regions of the tissue as it is stimulated with an electric field.

Importantly, the different edge guidance cues in the model of Equations (11)-(16) explicitly depend on the location of the unit normal at these fixed tissue locations. In a tissue of arbitrary shape, however, we would expect the normal to vary considerably along the edge. Moreover, the model of Equations (11)-(16) cannot explicitly describe the velocity at arbitrary tissue locations not contained in the discrete tissue regions it was designed for. For this reason, and since we are interested in controlling the velocity across the entire monolayer during electrotaxis, there is a need to scale up to a model that can describe the velocity of electrotaxis throughout the whole tissue. This will be the focus of the next section.

## 4 Construction and validation of a continuum model for collective electrotaxis

In the previous section, we studied spatial heterogeneity in collective electrotaxis by modelling the velocities of distinct regions in the tissue (*i*.*e*., the bulk and the edges). However, if one wishes to describe spatial heterogeneity during collective electrotaxis, it is important to understand how the velocity field can be described throughout the tissue, regardless of tissue geometry. The need to develop a model that works for arbitrary tissue geometries is further motivated by the fact that in practical applications of electrotaxis, one is interested in moving the tissue along arbitrary, and user-defined, trajectories in time and in space. For this reason, understanding how the entire monolayer moves is of the greatest interest in controlling collective electrotaxis in epithelial monolayers. Hence, we will focus in this section on developing and validating a continuum model for collective migration during electrotaxis that describes the evolution of monolayer velocities. Using the coupled system of Equations (11)-(16), we found that the behaviour of the edges during electrotaxis can be described as the result of reduced sensitivity to the electric field together with linear superposition of the cue to migrate outward at the edges. In this section, we seek to use these insights to formulate and analyse a continuum model for cell migration.

### 4.1 Model development

The goal of this section is to formulate a continuum model for the evolution of cell density during electrotaxis using a reaction-diffusion equation. At a continuum level, the migration of a population of cells is commonly modelled using a PDE for cell density, *ρ*, of the form

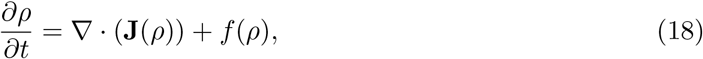

where **J** represents the *flux* of cells, and *f* is a function that represents cell proliferation [17]. Continuum models of cell migration are widely used [17] and provide physically or phenomenologically derived expressions for the fluxes and proliferation terms in Equation (18). For instance, in the context of chemotaxis, the standard Keller-Segel model decomposes the flux into a diffusive term due to random motion and one due to chemoattraction, *i*.*e*.,

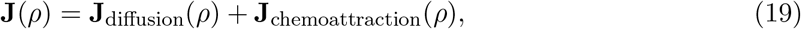

and a wide range of models exists to describe the different (nonlinear) diffusion and chemoattraction terms. In this work, we will assume the flux due to random movement of cells to be given by non-linear diffusion of the form

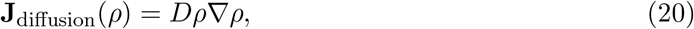

where *D >* 0 is the diffusion coefficient. This model has been previously shown to well describe the edge spreading of epithelia [18]. We also choose the proliferation term to be described by the standard logistic growth model, which is a canonical model for cell proliferation and takes the form

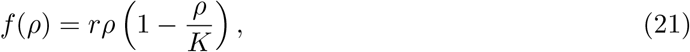

where *r >* 0 is the proliferation rate, and *K >* 0 is the carrying capacity [17]. Having fixed the flux due to random cell movement and the proliferation function, we turn to proposing a model for the tissue flux due to electrotaxis, which is the main contribution of this paper. In his seminal work [17], Murray, drawing an analogy with chemotaxis, proposed that the flux due to electrotaxis be represented as

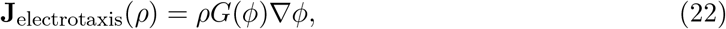

where *ϕ* is an electric potential, and *G* is a function that encodes the strength of the response of cells to the electric field. This model assumes that the response to the electric field is in the direction of the gradient of the potential, and that the strength of the response is dependent on the local magnitude of the potential. While the first assumption is reasonable, given the experimental evidence presented in this paper, the preceding analysis demonstrating that the tissue edges are less sensitive to the electric field gives an indication that edge-effects must be included in a continuum model for collective electrotaxis. In a continuum model describing tissue density, the difference in density at the edge versus in the bulk gives a good proxy to quantitatively establish the location of the tissue edge. For this reason, we incorporate the spatial heterogeneity in cell responses to the electric field by choosing a functional form for *G* that makes responses to the electric field strongest in the bulk of the tissue, where *ρ* ≈ *K*, and weakest at the boundary where *ρ* ≈ 0. Since calibrating the system of Equations (11)-(16) shows that the ratio of the sensitivity at the edges to that in the bulk can be described by a fixed parameter, *δ* ∈ [0, 1], a simple model for the dependence of the electrotaxis flux on local cell density is given by

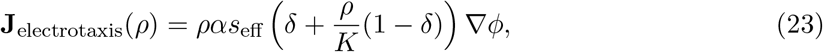

and *α >* 0 represents the strength of the electric field response in the bulk, such that

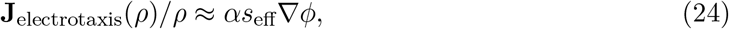

in the tissue bulk and

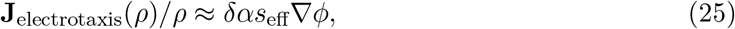

at the tissue edges, which corresponds with the analysis in the Bayesian inference of Equations (11)-(16). We note here that *s*_eff_ i s n ow a s patially varying fi eld. Br inging to gether the fluxes and the proliferation term, we propose the following continuum model for collective electrotaxis,

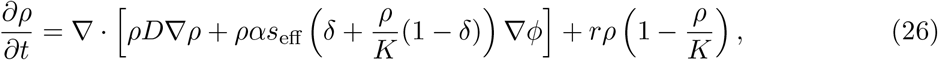

subject to an initial condition describing the cell density at the beginning of electrotaxis and boundary conditions for the domain. In this paper, we consider initial conditions corresponding to the cell density at *t* = 0h of the experiment, and we consider no-flux boundary conditions for the computational domain that contains the cell monolayer. So far, for the evolution of the effective signal, we have used the governing equations in Equations (11)-(16), since the electric field was assumed to be uniaxial. In a fully two-dimensional model, however, the electric field can locally point in any arbitrary direction, and its magnitude can also vary in space. For that reason, we must define an extension for the governing equations for the effective signal, *s*_eff_, that takes into account a vector field for the velocity, whose magnitude can vary in space. Here, we make the assumption that the strength of the effective signal only depends on the magnitude of the electric field, and so rewrite the governing equations for the effective signal and the inhibitor

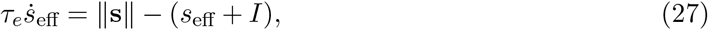

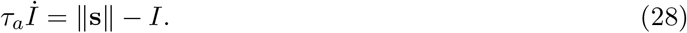

To conclude, we have proposed Equations (26)-(28) as a model extension to the model in Equations (1) and (2) that takes into account several edge effects at the tissue boundaries. This model contains three additional parameters: the strength of the response to the electric field when the preferred polarity of the tissue is perpendicular to the electric field, *α*_⊥_; the difference in field responsiveness, *δ*; and the strength of the endogenous cue, *α*_edge_.

### 4.2 Bayesian inference of model parameters

We wish to calibrate the model in Equations (26)-(28) to the experimental cell density data of Wolf *et al*. [7]. However, fitting a full two-dimensional density model to the data is challenging due to the very large local fluctuations in cell density, which are characteristic of MDCK epithelial monolayers during expansion [19,20]. For this reason, we select a region consisting of a horizontal strip of the monolayer which is one mm away from the top and bottom edges and average the cell density in the *y*-direction. This averaging removes any artefacts that arise from migration from the top and bottom edges and provides an averaged one-dimensional density profile across the monolayer. In one dimension, Equation (26) reduces to

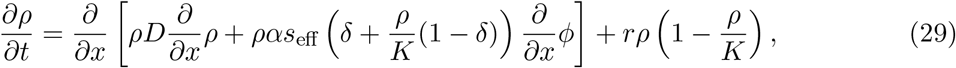

subject to initial conditions given by the experimentally measured tissue density at *t* = 0h, and Neumann boundary conditions on the right and left boundaries of the domain given by the field of view containing the monolayer in Figure 1. The domain is the interval spanned by the *x*-coordinate in the field of view of the experiment shown in Figure 1, *i*.*e*., the range -3.5mm to 3.5mm from the tissue centre. The full parameter set, *θ*, for the model in Equations (27)-(29) is given by

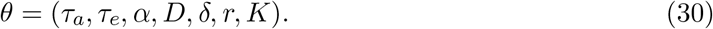

In the experiments of Wolf *et al*. [7], the field was uniaxial and constant in space and time, meaning that the temporal dynamics of the effective signal, *s*_eff_, are homogeneous in time and are identical to those described in Section 3. For this reason, since the parameters governing the temporal dynamics of the effective signal, *τ*_*a*_ and *τ*_*e*_, could be confidently identified given the data, we choose the posterior means for *τ*_*a*_ = 2.04h and *τ*_*e*_ = 0.26h from Section 3 and perform Bayesian inference on the remaining model parameters, *α, D, δ, r*, and *K*.

Using the same Bayesian inference procedure as in Section 3, we obtain the posterior distributions for the model parameters *α, D, δ, r*, and *K*, with their posterior means recorded in Table 2. As posterior means we choose uniform distributions for *α* ∈ [0, 2]mm/h, *D* ∈ [0, 20]mm^2^h^−1^, *δ* ∈ [0, 1], *r* ∈ [0, 1]h^−1^, and *K* ∈ [0, 300]mm^−1^. The marginal posterior distributions of the model parameters are shown in Figure 3; all model parameters can be confidently estimated given the experimental data with each of the distributions supported on a narrow interval around its mean. To understand the quality of the resulting model predictions given the posterior distributions over model parameters, we simulate from the model in Equations (27)-(29) using the posterior means of the distributions shown in Figure 3, and the initial condition from the data as the initial condition in the simulations. Figure 4 shows a comparison between the experimental data and the predictions generated by the model. The kymographs in Figure 4 show a density profile across the tissue that is in excellent agreement with the experimental data. Figure 4 also shows the profiles of the leading and the trailing edge at one, two, and three hours during electrotaxis as simulated by the model, compared to the experimental data, again showing excellent agreement between the model predictions and the experimental data. We conclude that the model in Equations (27)-(29) can faithfully capture the migratory behaviours of the monolayer during electrotaxis, and can be used to predict cell densities during collective migration when the tissue is subject to an electric field.

**Table 2:** Posterior means for the extended edge velocities model. Note that the value for the velocity decay rate, *γ*, is estimated directly from the data.

| $\alpha$ (mm/h) | $D$ (mm<sup>2</sup>h<sup>-1</sup>) | $\delta$ | $r$ (h<sup>-1</sup>) | $K$ (mm<sup>-1</sup>) |
| --- | --- | --- | --- | --- |
| 0.837 | 13.30 | 0.625 | 0.180 | 260.29 |

**Figure 3:**
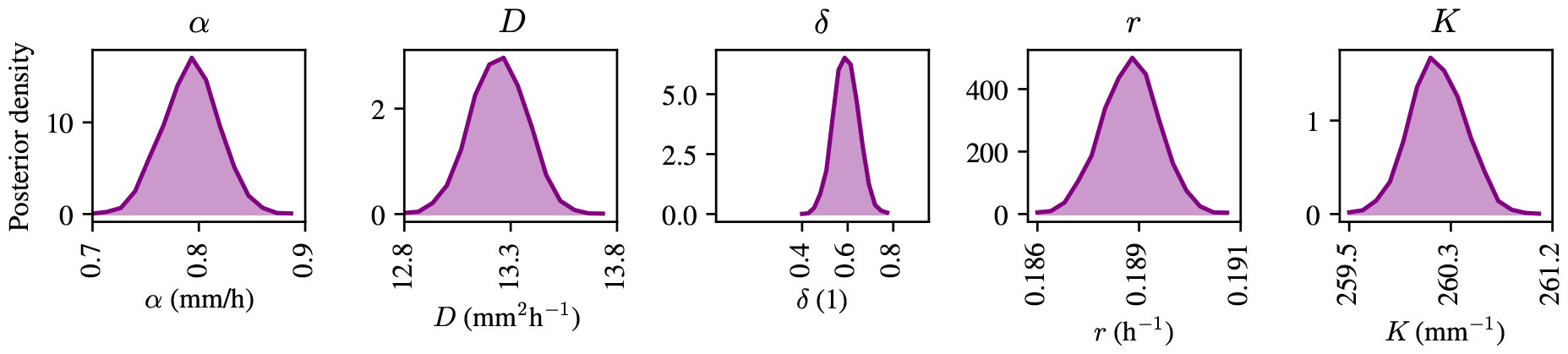
Marginal posterior distributions for a one-dimensional continuum model of collective electrotaxis in Equations (27)-(29).

**Figure 4:**
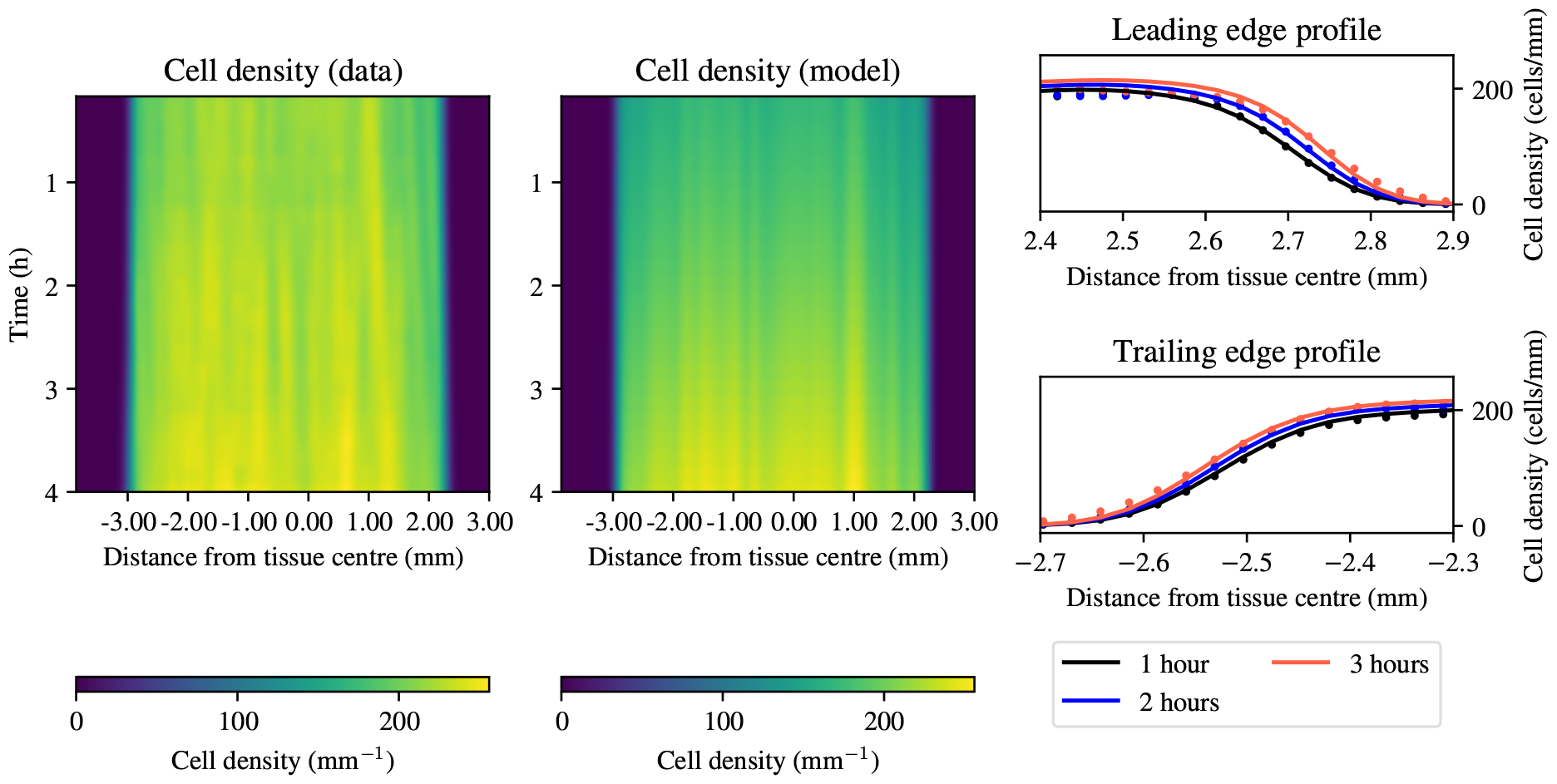
Comparison between model predictions and experimental data of one-dimensional cell density profiles during collective electrotaxis. Left panels: kymographs for cell density (experimental data left, model solutions right). Right panel: profiles of the leading and trailing edge at one, two, and three hours of electrotaxis.

### 4.3 Tissue size affects migratory speed

The model in Equations (27)-(29) provides a good description of the evolution of cell densities during electrotaxis. Now, we wish to return to the two-dimensional case and outline in this section how the model in Equation (26) can be used to obtain predictions for the local migratory velocity. We will use this framework to show how tissue size and geometry affect the magnitude, and distribution, of migratory velocity in MDCK monolayers. Inspection of Equation (26) shows that the predicted velocity of migration is given by

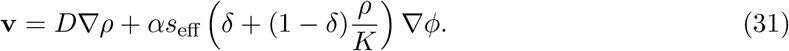

The expression in Equation (31) allows us to compute the velocity field from simulations of the model in Equation (26), and it allows us to probe the role of tissue geometry and size in the resulting tissue velocities during collective electrotaxis.

To study the velocity of the monolayer during electrotaxis, we simulate tissues with two different geometries: circular and triangular. These shapes provide a good starting point for understanding the role of tissue geometry, given that they have curvature and acute corners. For each shape, we simulate tissues with a diameter between 0.25mm and 4mm, corresponding roughly to the range between the smallest tissue sizes in which directional collective electrotaxis has been reported [21] and twice the size of the experiments of Wolf *et al*. [7], thus providing a good range of realistic tissue sizes for collective electrotaxis. For each tissue shape, we generate a grid of tissue diameters consisting of 100 equally spaced points between this minimum and maximum tissue diameter, and compute the maximum electrotaxis velocity in the simulation, using the means of the posterior distributions of the model parameters obtained in Figure 3. For all model simulations, a uniform initial condition of *ρ* = *K* within the tissue is used. We solve Equation (26) numerically in two spatial dimensions with the finite-volume numerical scheme described in [18, 22].

Figure 5 shows that tissue size has an impact on the maximum velocity during electrotaxis: with increasing tissue size, the maximum velocity during electrotaxis increases before reaching a plateau. The scales of the variation in tissue velocity are vastly different between the two different geometries, as shown by the limits of the *y*-axes in Figure 5. The difference in the maximum velocity between the smallest triangular tissues and the largest triangular tissues amounts to less than 0.5*μ*mh^−1^, whereas it is more than 10*μ*mh^−1^. The insets in Figure 5 show that the spatial distribution of velocities is qualitatively different in the circular tissues of different sizes, with the large circular tissue and both triangular tissues having a well-defined edge with a lower magnitude velocity. Intuitively, this shows that the locations and orientations of the tissue edges have an effect on the migratory velocity of the tissue as a whole, and that these effects must be taken into account when designing bespoke electric fields to achieve collective migration in tissues with different shapes. In summary, we find that tissue geometry geometry, together with tissue size, informs the distribution, and magnitude of the velocity in the electrotaxing monolayer.

**Figure 5:**
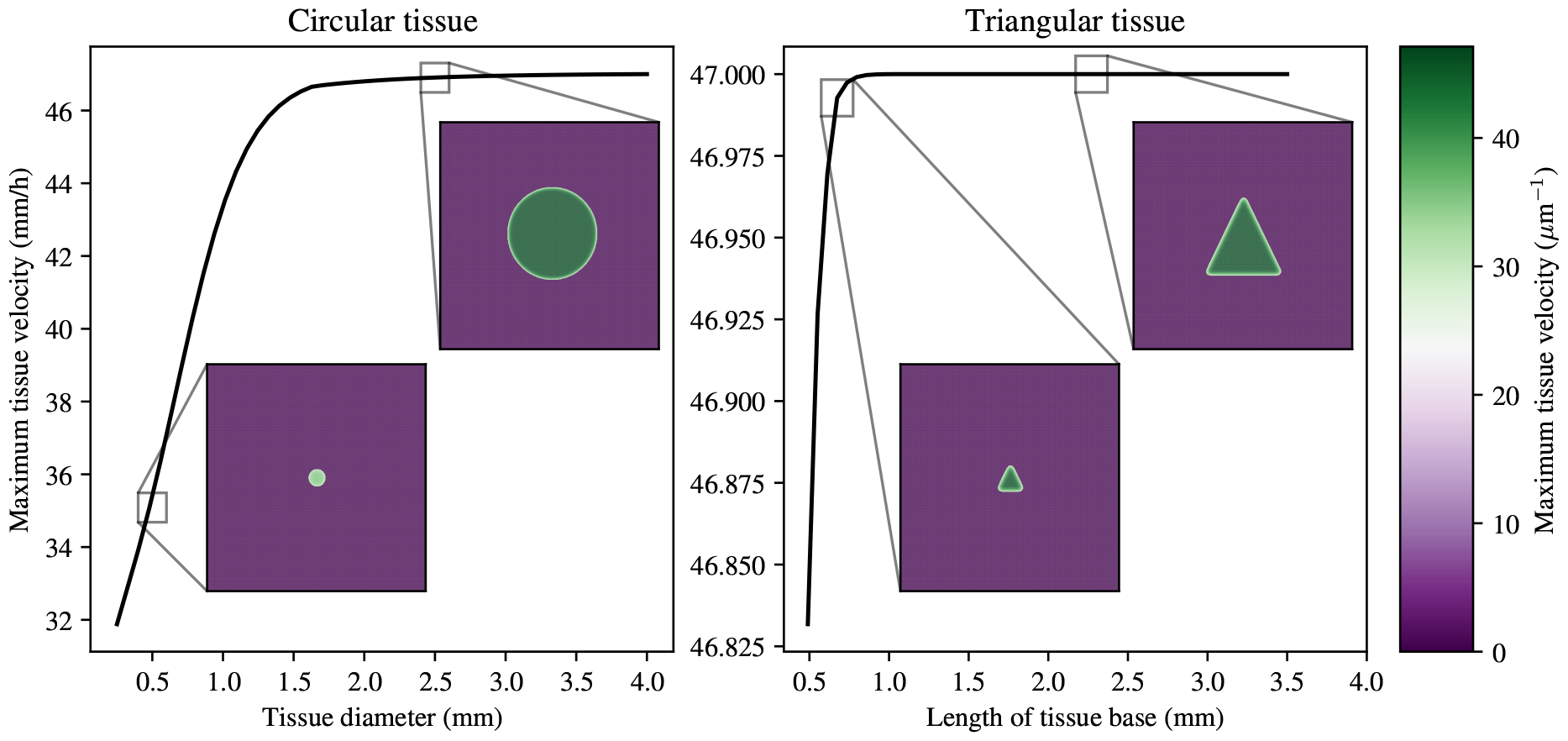
Effect of tissue size and geometry on maximum migratory velocity during electrotaxis. Insets show circular (triangular) tissues with a tissue base of 0.5mm and 2.5mm, respectively. The spatial distribution of velocities is qualitatively different in the circular tissues, but not in the triangular tissues.

## 5 Optimal design of electric fields to achieve spatially uniform migration velocity

Having understood that the continuum model from Equation (26) provides an expression for the tissue velocity, **v**, through Equation (31), we can turn to the question of how to use these insights to design bespoke electric fields to achieve a target spatial velocity distribution during collective electrotaxis. While the possible applications of electrotaxis to guide collective cell migration are myriad [3], we focus on one specific task in controlling collective migration, which is to create a spatially uniform migratory velocity. Our motivation for this is that overcoming the large differences in migration velocities between the edges and the bulk of the tissue, which are a result of the interplay between the endogenous and exogenous cues that cells at different tissue locations receive, would represent an important first example of spatial control of collective velocity during electrotaxis. In this section, we first investigate the spatial distribution of an optimal electric field predicted by a one-dimensional model, which can be analysed analytically, and conclude with two numerical examples in two dimensions to demonstrate practical applications of the method.

### 5.1 Physical insights from a one-dimensional model

Recall that in one dimension, the electrotaxis model for the evolution of cell densities is given by Equations (27)-(29), and the velocity can be found analogously to that in Equation (31),

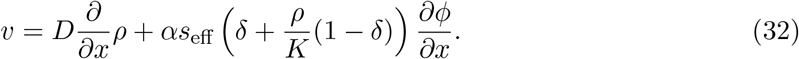

We now turn to the following problem. Given a target velocity, *v*^⋆^ *>* 0, design an electric potential, *ϕ*, such that the monolayer velocity is uniform, *v* = *v*^⋆^. When *s*_eff_ *>* 0, Equation (32) rearranges to

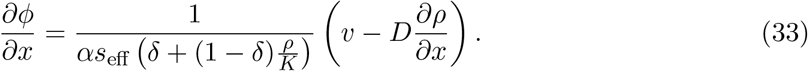

While tractable, the expression for the field in Equation (33) comes with the limitation that it depends on *s*_eff_, which is zero at *t* = 0. As we have discussed previously, *s*_eff_ also exhibits *s*_eff_ → 0 as *t* → ∞. Therefore, Equation (33) does not prescribe how to start stimulation at *t* = 0, and predicts unphysiologically strong electric potential gradients as *s*_eff_ → 0 for large *t*. For this reason, the expression for the field in Equation (33) does not constitute a full optimal control solution. At this point, we also note that controlling the velocity, *v*, in Equation (33) so that it matches some desired velocity is not a straightforward problem, conceptually, or computationally due to the highly nonlinear coupling with the effective signal, *s*_eff_. This is made more complicated by the fact that in applications more generally, one might want the target velocity, *v*^⋆^, to vary in space. For that reason, we make progress by simplifying the problem substantially. Instead of considering a full optimal control problem, we propose to overcome the difficulty that *s*_eff_ = 0 at *t* = 0 by stimulating with a constant electric field until *s*_eff_ is at a sufficient level that the gradient in Equation (39) can be used. As *s*_eff_ wanes, we propose that the electric potential gradient is capped by a maximal field strength, *i*.*e*.,

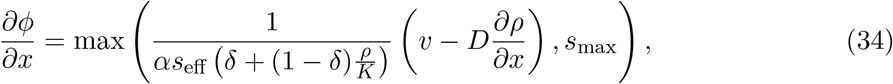

for some *s*_max_ *>* 0 that describes the maximum field strength that is physiologically tolerable. With this in mind, we can turn to understanding how the one-dimensional model of electrotaxis in Equations (27)-(29) can be used to obtain electric fields to achieve a constant velocity of migration throughout the monolayer. By substituting *v* = *v*^⋆^ for the velocity and integrating over the domain, one obtains the following direct expression for the potential,

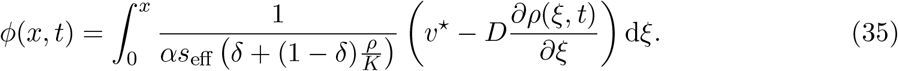

Here, the lower limit of integration can be chosen arbitrarily, since the velocity in Equation (32) is only influenced by the gradient of the potential, ∂_*x*_*ϕ*, meaning that the constant of integration is arbitrary. This also is equivalent to the natural boundary condition

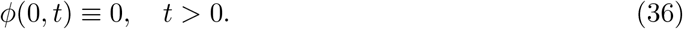

With this in mind, the problem of keeping the velocity constant in the monolayer reduces to

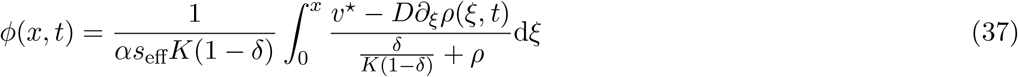

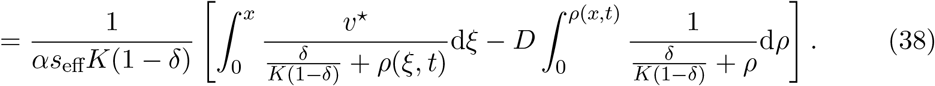

Evaluating the final integral yields

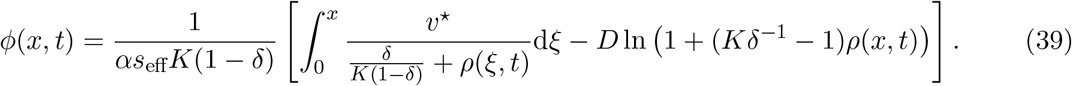

The two terms in the potential defined in Equation (39) carry convenient physical interpretations. The first term in Equation (39) describes the changes in potential needed for a tissue element at position *x* to migrate at the same velocity as the trailing edge. The second term in Equation (39) corresponds to corrections in the field strength that arise from differences in the sensitivity of cells that are at the bulk or at the edge of the tissue.

### 5.2 Design of electric fields for arbitrary shapes

In two dimensions, the coupling of velocity to local gradients in the electric potential, *ϕ*, and cell density can also be obtained explicitly from the equation for the tissue velocity, Equation (31). As in the previous section, one can set the problem of designing an electric potential, in this case in two dimensions, such that the entire tissue is moving at the same velocity. We denote this velocity by **v**^⋆^. In a more general optimal control framework, the target velocity of migration, **v**^⋆^, could vary in time and in space. In the present framework, whenever *s*_eff_ *>* 0, Equation (31) stipulates that the optimal gradient of the electric potential, which we denote by **s**_⋆_, is given by

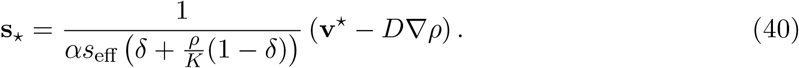

As in the one-dimensional case discussed in Section 5.1, there are limits on the velocities that can be physically achieved, as the system adapts to the electric field signal: Equation (40) fails to hold when *s*_eff_ = 0 and predicts unphysically strong potential gradients in the limit as *s*_eff_ → 0. At the same time, having a spatially varying field, **s**_⋆_, makes the effective signal, *s*_eff_, vary in space, thereby making the computation of the optimal field through Equation (40) significantly more complicated. Therefore, to avoid the complications that could arise from a spatially varying *s*_eff_ on the one hand, and possibly unphysically strong electric fields on the other, we propose a simple heuristic to achieve migration in the direction of the desired velocity. That is, we let the direction of the field vary, but not its magnitude. This comes with the limitation that while the resulting field will be in the direction of the optimal field predicted by Equation (40), the magnitude and the temporal distribution might not be optimal for the desired target velocity. We suggest this problem as a starting point for future research. For now, in order to make progress in the design of two-dimensional electric fields for controlling collective electrotaxis, the heuristic we propose is to choose a fixed magnitude for the electric field, in the direction of the optimal gradient, *s*_⋆_, as defined in Equation (40). Given a fixed strength of the field, *s*_max_, this amounts to setting the gradient of the potential, ∇*ϕ*, as

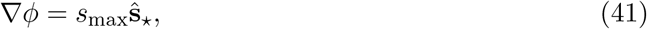

where 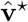 ^⋆^, is a unit vector in the direction of the optimal field, **s**_⋆_. This heuristic electric field provides a means to vary the electric field in space to effectively counteract the influence of local gradients in tissue density, which can cause the tissue to spread in directions other than that of the target velocity. As a proof of principle, we take as target velocity, **v**^⋆^, a constant vector field in the *x*-direction with constant magnitude given by the maximum velocity of the bulk in Figure 1, *i*.*e*., 47*μ*mh^−1^. We take two sample geometries, one circular with a diameter of 3.5mm, and one triangular with a tissue base of 3mm, initialised using uniform initial condition of *ρ* = *K*, before applying the electric field suggested by Equation (41). After having observed that the field is near constant in time, we plot the gradient, **s**_⋆_, in Figure 6.

**Figure 6:**
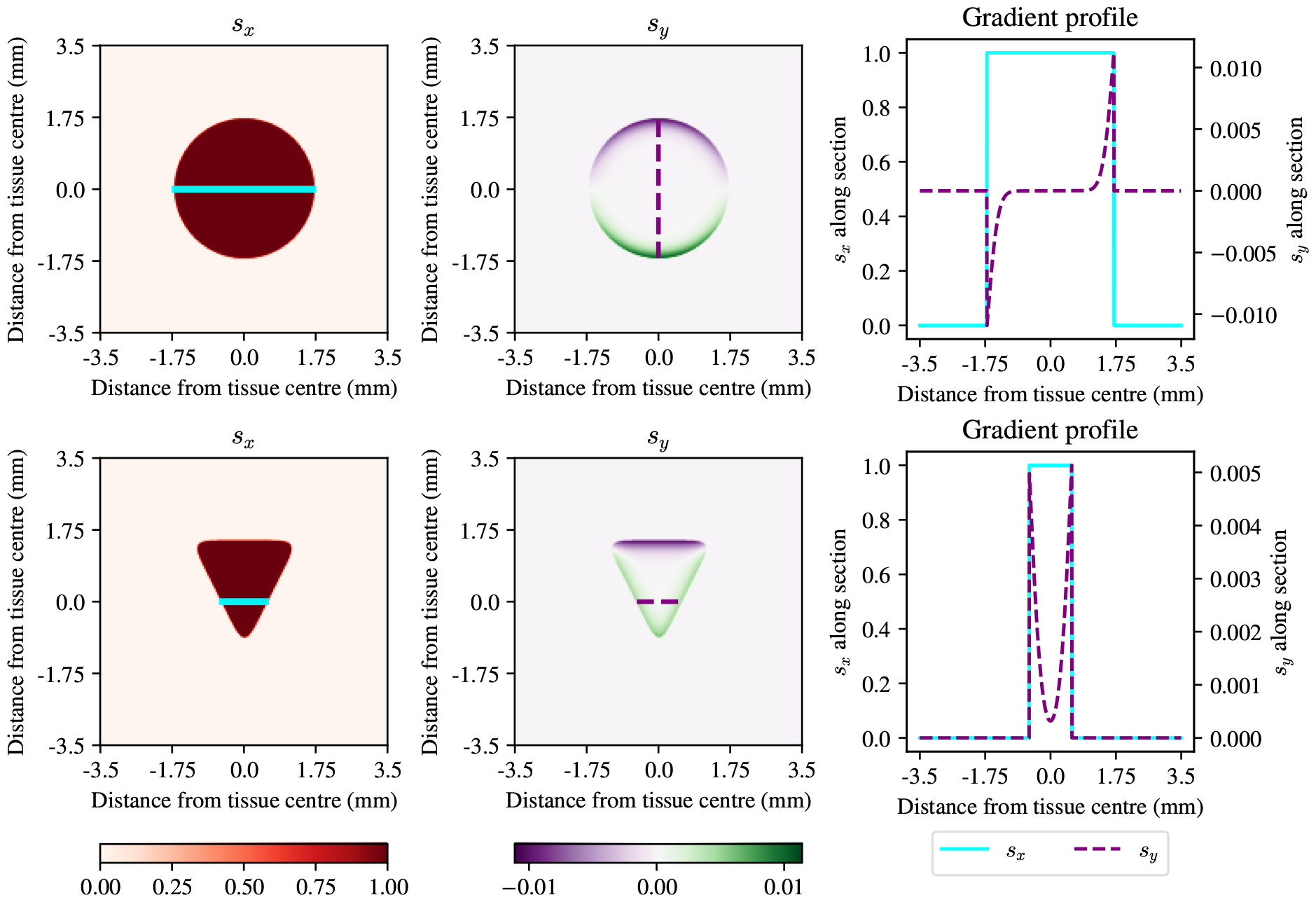
Spatial distribution of the electric field after one hour of electrotaxis to achieve uniform migration in the *x*-direction, for a circular tissue (top row) and a triangular tissue (bottom row). First column: *x*-component of the heuristic electric field. Second column: *y*-component of the heuristic electric field. Third column: one-dimensional profiles along solid cyan and dashed purple lines in the two-dimensional tissues, respectively.

Figure 6 shows that the electric field computed using the heuristic in Equation (41) mainly acts to override the outgrowth of the epithelium in directions other than the electric field and translate the tissue in the *x*-direction. For example, in the circular tissue there is a clear edge region defined by having a *y*-component of opposite sign to that of the unit normal at the tissue boundary, with a similar phenomenon occuring in the triangular tissue. The one-dimensional profiles in Figure 6 show that the gradient acts against the outward migration of the epithelial monolayer in a small region at the tissue boundary.

## 6 Discussion

In this paper, we have investigated the mechanisms behind the spatial heterogeneity in collective electrotaxis that was first observed in the experiments of Wolf *et al*. [7], and we have used these insights to develop a continuum model of collective electrotaxis. This model lends itself to the experimental design of bespoke electric fields that can be used to achieve specific aims during engineered collective cell migration. There are several ways in which our modelling approach can be enhanced for applications in bio-engineering on the one hand, and for a better understanding of the cell signalling and mechanics involved in establishing collective electrotaxis, on the other.

The specific bioengineering application of the continuum model proposed in this work consisted of the design of an electric field to achieve uniform tissue velocity across the tissue. However, our model can equally be applied in situations in which the desired velocity field is not spatially uniform, but varies in space. Such applications exist, for example, in wound healing, where one would want to move two opposite sides of a wound closer together to achieve healing. A key area for future research is then the construction of a full optimal control formulation for spatially and temporally varying electric fields. Formulating the design of spatially varying electric fields as an optimal control problem would overcome two restrictions that we introduced with the heuristic in Equation (41). First, an optimal control formulation would cast the design of an electric field as minimising a given objective function that can encode the desired monolayer displacement trajectory during electrotaxis. This allows for a huge range of possible electric field designs and applications. In practice, there exist many examples of spatially varying fields to induce electrotaxis [1, 3, 23]. Therefore, the ability to implement bespoke spatially varying electric fields can be exploited to directly implement the predictions for an optimal electric field in practical applications. Second, having a coupled system of PDEs describing the optimal electric field would provide a stimulation protocol starting from *t* = 0 where *s*_eff_ = 0. This would avoid having to stimulate with an arbitrarily set field first to make sure that *s*_eff_ is large enough to use the field in Equation (40).

The second, and final area of extension for this work concerns the phenomenon of undershoot at the top and trailing edges seen in Figure 1. While the results of Section 3.1 show that the system of ODEs given in Equations (11)-(16) can accurately represent the velocities of different tissue locations during electrotaxis, one of the phenomena in the data in Figure 1 that is not well-explained by the model is the fact that the top and trailing edges display an undershoot of the velocity when the field is switched off. We explore several possible starting points for the modelling of this phenomenon in Supplementary Information Section S2. We show that while mechanical factors seem unlikely to contribute to the undershoot at both the top and the trailing edge, intracellular signalling could potentially explain the undershoot observed on long time scales at the top edge of the tissue only. In this view, the effective signal becomes negative when the electric field is turned off, indicating that the internal signalling pathways in the constituent cells transduce a signal to reverse cell polarity. This phenomenological modelling finding begs the question of how such an intracellular signal can come about, and how it should be explored in future efforts. In this regard, we believe an interdisciplinary approach combining model predictions and experimental implementations can shed light on the dynamics involved: for example, model-based predictions of collective migration under different electric field strengths and temporal dynamics can be directly compared with experimental data of collectively electrotaxing epithelia under such electric fields. This would allow us to test the validity and applicability of such models in predicting the response of cells in different tissue regions, thus providing a more detailed mechanistic understanding of the temporal signalling dynamics involved.

## Authors’ contributions

S.M.P. conceived the project, developed the mathematical modelling, created the code for the numerical implementation, produced all figures, and carried out the numerical experiments and analysis; R.E.B. helped design, supervised and coordinated the study. S.M.P. wrote the paper, on which R.E.B., D.J.C., and I.B.B. commented and revised. All authors gave final approval for publication.

## Acknowledgements

S.M.P. is supported by an EPSRC/UKRI Doctoral Training Award. I.B.B. is supported by an NSF GRFP. D.J.C. would like to acknowledge support for this work was provided in part by the National Institute of Health Award R35 GM133574-03 and the National Science Foundation CAREER Award 2046977. R.E.B. was supported by a grant from the Simons Foundation (MP-SIP-00001828).

## References

[1] Daniel J. Cohen, W. James Nelson, and Michel M. Maharbiz. Galvanotactic control of collective cell migration in epithelial monolayers. Nature Materials, 13(4):409–417, 2014.

[2] Greg M. Allen, Alex Mogilner, and Julie A. Theriot. Electrophoresis of cellular membrane components creates the directional cue guiding keratocyte galvanotaxis. Current Biology, 23(7):560–568, 4 2013.

[3] Tom J Zajdel, Gawoon Shim, Linus Wang, Alejandro Rossello-Martinez, and Daniel J Cohen. SCHEEPDOG: programming electric cues to dynamically herd large-scale cell migration. Cell systems, 10(6):506–514, 2020.

[4] Simon F Martina-Perez, Isaac B Breinyn, Daniel J Cohen, and Ruth E Baker. Optimal control of collective electrotaxis in epithelial monolayers. arXiv preprint 2402.08700, 2024.

[5] C. Blanch-Mercader, R. Vincent, E. Bazellières, X. Serra-Picamal, X. Trepat, and J. Casademunt. Effective viscosity and dynamics of spreading epithelia: a solvable model. Soft Matter, 13(6):1235–1243, 2017.

[6] Ricard Alert and Xavier Trepat. Physical models of collective cell migration. Annual Review of Condensed Matter Physics, 11(1):77–101, 2020.

[7] Abraham E Wolf, Matthew A Heinrich, Isaac B Breinyn, Tom J Zajdel, and Daniel J Cohen. Short-term bioelectric stimulation of collective cell migration in tissues reprograms long-term supracellular dynamics. PNAS Nexus, 1(1), 3 2022.

[8] Yan Zhang, Guoqing Xu, Jiandong Wu, Rachel M Lee, Zijie Zhu, Yaohui Sun, Kan Zhu, Wolfgang Losert, Simon Liao, Gong Zhang, Tingrui Pan, Zhengping Xu, Francis Lin, and Min Zhao. Propagation dynamics of electrotactic motility in large epithelial cell sheets. iScience, 25(10):105136, 2022.

[9] Radek Erban and Hans G. Othmer. From signal transduction to spatial pattern formation in E. coli: A paradigm for multiscale modeling in biology. Multiscale Modeling and Simulation, 3(2):362–394, 2005.

[10] Shreyansh Jain, Victoire M. L. Cachoux, Gautham H. N. S. Narayana, Simon de Beco, Joseph D’Alessandro, Victor Cellerin, Tianchi Chen, Mélina L. Heuzé, Philippe Marcq, René Marc Mège, Alexandre J. Kabla, Chwee Teck Lim, and Benoit Ladoux. The role of single-cell mechanical behaviour and polarity in driving collective cell migration. Nature Physics, 16(7):802–809, 7 2020.

[11] Satomi Matsuoka and Masahiro Ueda. Mutual inhibition between PTEN and PIP3 generates bistability for polarity in motile cells. Nature Communications, 9(1), 12 2018.

[12] Changji Shi, Chuan Hsiang Huang, Peter N. Devreotes, and Pablo A. Iglesias. Interaction of motility, directional sensing, and polarity modules recreates the behaviors of chemotaxing cells. PLoS Computational Biology, 9(7), 7 2013.

[13] Gillian DeWane, Alicia M. Salvi, and Kris A. DeMali. Fueling the cytoskeleton-links between cell metabolism and actin remodeling, 2 2021.

[14] Sham Tlili, Estelle Gauquelin, Brigitte Li, Olivier Cardoso, Benoît Ladoux, Hélène Delanoë Ayari, and Francois Graner. Collective cell migration without proliferation: Density determines cell velocity and wave velocity. Royal Society Open Science, 5(5), 5 2018.

[15] Michael Clerx, Martin Robinson, Ben Lambert, Chon Lok Lei, Sanmitra Ghosh, Gary R. Mirams, and David J. Gavaghan. Probabilistic Inference on Noisy Time Series (PINTS). Journal of Open Research Software, 7:1–6, 2019.

[16] Aki Vehtarh, Andrew Gelman, Daniel Simpson, Bob Carpenter, and Paul Christian Burkner. Rank-Normalization, folding, and localization: an improved (formula presented) for assessing convergence of MCMC (with discussion). Bayesian Analysis, 16(2):667–718, 2021.

[17] James D. Murray. Mathematical Biology I : an introduction. Interdisciplinary applied mathematics; 17. Springer, New York, N.Y, 3rd ed. edition, 2002.

[18] Carles Falcó, Daniel J. Cohen, Jose A. Carrillo, and Ruth E. Baker. Quantifying tissue growth, shape and collision via continuum models and Bayesian inference. Journal of the Royal Society Interface, 20(204), 2023.

[19] Shao-Zhen Lin, Peng-Cheng Chen, Liu-Yuan Guan, Yue Shao, Yu-Kun Hao, Qunyang Li, Bo Li, David A Weitz, and Xi-Qiao Feng. Universal statistical laws for the velocities of collective migrating cells. Advanced Biosystems, 4(8):2000065, 2020.

[20] Lin Shao-Zhen, Wu-Yang Zhang, Dapeng Bi, Bo Li, and Xi-Qiao Feng. Energetics of mesoscale cell turbulence in two-dimensional monolayers. Communications Physics, 4(1):21, 2 2021.

[21] Li Li, Robert Hartley, Bjoern Reiss, Yaohui Sun, Jin Pu, Dan Wu, Francis Lin, Trung Hoang, Soichiro Yamada, Jianxin Jiang, and others. E-cadherin plays an essential role in collective directional migration of large epithelial sheets. Cellular and Molecular Life Sciences, 69(16):2779–2789, 2012.

[22] Rafael Bailo, José A. Carrillo, Hideki Murakawa, and Markus Schmidtchen. Convergence of a fully discrete and energy-dissipating finite-volume scheme for aggregation-diffusion equations. Mathematical Models and Methods in Applied Sciences, 30(13):2487–2522, 12 2020.

[23] José Leal, Sebastian Shaner, Nicole Jedrusik, Anna Savelyeva, and Maria Asplund. Electrotaxis evokes directional separation of co-cultured keratinocytes and fibroblasts. Scientific Reports, 13(1):11444, 7 2023.

