## Supplementary Information for "Spatial heterogeneity in collective electrotaxis: continuum modelling and applications to optimal control"

Simon F. Martina-Perez, Isaac B. Breinyn, Daniel J. Cohen, and Ruth E. Baker

#### S1 Estimating the velocity decay rate from the data directly

In this section, we detail the estimation of the velocity decay parameter, $\gamma$, directly from the data. We have done this estimation previously on the same dataset from Wolf *et al.* [1] in our previous work [2] and report it here for completeness. As noted in the main text, the constituent equations relating the effective signal, $s_{\text{eff}}$, to the velocity predict that, given an initial velocity, $v_0$, the velocity in the bulk of the tissue decays exponentially in the absence of an electric field. Since the experimental data contain information on the bulk velocity after stimulation with the electric field is turned off, it follows that the observed decaying velocity profile can be used to immediately infer the decay rate, $\gamma$. From these considerations, we seek to fit an exponential of the form

$$v(t) = Ce^{-\gamma(t-t_{\text{end}})}, \tag{S1}$$

to the bulk velocity data in the 2.5 hours after stimulation with the electric field is turned off. Here, $t_{\text{end}}$ is the time at which the electric field is switched off, and $C$ is given by

$$C = v(t_{\text{end}}). \tag{S2}$$

We perform a least-squares fit for the decay rate, $\gamma$, to give an estimate of $\gamma = 1.765h^{-1}$. Figure A shows that the fitted exponential decay curve is in excellent agreement with the experimental data after pulse stimulation. This motivates us to take the value of $\gamma$ as fixed going forward. The benefit of directly estimating $\gamma$ from data is that it reduces the dimensionality of the parameter space, thus simplifying inference of the remaining parameters.

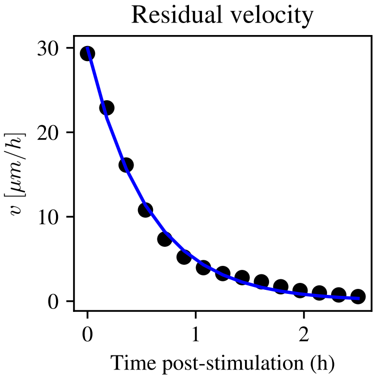


Figure A: Experimental data of bulk velocity decay post-stimulation with an electric field pulse of 3 V/cm (black dots) together with least-squares fit of an exponential decay curve in the form $C\exp(-\gamma t)$ (solid blue line). The least-squares solution gives an excellent fit to the data and can be used to estimate the decay rate, $\gamma$.

#### S2 Understanding recoil in the top edge post-stimulation

Here, we turn attention to understanding the phenomenon that the velocity in the top edge of the tissue becomes negative for several hours after the electric field is switched off. This finding was originally reported by Wolf *et al.* [1] who refer to it as recoil. In fact, recoil was also mentioned by Wolf *et al.* for the velocity at the trailing edge, which exhibits a sharp drop below its equilibrium value before restoring. On this phenomenon, Wolf *et al.* proposed that the observed recoil might be due to a mechanical effect induced by the visco-elastic properties of the epithelium. Although such a mechanical effect may indeed be at play at the trailing edge, we reason that it is not sufficient to explain the sustained negative velocity at the top tissue edge, as the characteristic time scale for the elastic response of MDCK epithelia is on the order of 15 to 30 minutes [3]. Therefore, the recoil as observed in the experiments of Wolf *et al.* [1] cannot be explained by visco-elastic mechanical effects alone in the top edge. We then ask the question: if mechanics are not responsible for creating recoil, might it be that cessation of the electric field signal itself could be inducing cells to move in the opposite direction of the electric field post-stimulation?

##### S2.1 Model and constitutive equations

So far, we have assumed that the effective signal inside the constituent cells making up the monolayer is always positive given a positive field, as we have considered $s^{+}_{\text{eff}}$ as in Equation (17) in the main text. The effect of this is that the effective signal inside the cells can only give a migratory command in the direction of the electric field. To understand whether cessation of the electric field itself can provide a negative signal to the cells, we wonder if the model assumption that the effective signal is always in the direction of the field can be relaxed, *i.e.* by replacing $s^+_{\text{eff}}$ by $s_{\text{eff}}$ in the active force for the top edge. Since we only consider the dynamics of the top edge in this case, we can considerably simplify the model at the top edge, since there is no need to compare the response of the top edge relative to that of the bulk. Without loss of generality, we can set $\delta = 1$, and study a reduced system of equations describing the dynamics of the top edge of the tissue,

$$\dot{v}_{\text{top}} = -\gamma v_{\text{top}} + \alpha s_{\text{eff}}, \tag{S3}$$

$$\tau_e \dot{s}_{\text{eff}} = s - (I + s_{\text{eff}}), \tag{S4}$$

$$\tau_a \dot{I} = s - I. \tag{S5}$$

#### S2.2 Bayesian parameter identification

As before, we keep the pre-fit value for $\gamma$ constant, and seek to estimate the parameter values for $\alpha$, $\tau_e$, and $\tau_a$ using the same Bayesian inference procedure as in Section 3 of the main text. We find that the posterior means of the model parameters are given by 0.103h, 6.11h, and 57.37h$^{-1}$, respectively. By computing the posterior predictive interval of the middle 95% of model solutions generated using parameter values drawn from the posterior distribution, we observe in Figure B that the model in Equations (S3)-(S5) is able to accurately recapitulate the long-term undershoot of the average velocity in the top edge post-stimulation. Given this excellent agreement with the experimental data, we turn to understanding the quality of the posterior distributions over the model parameters.

The marginal posterior distributions over the model parameters $\alpha$, $\tau_a$, and $\tau_e$ in Figure B show that while the field response, $\alpha$, and the adaptation time scale, $\tau_a$, can be confidently identified from the data, the excitation time scale, $\tau_e$, is less confidently estimated. The marginal posterior distribution for the excitation timescale, $\tau_e$, is broad and largely uninformative. This is different from the posterior distribution for $\tau_e$ in Section 3 of the main text, where $\tau_e$ did not suffer this problem and could be confidently identified from the data. When doing inference on only the data of the top edge, while the posterior distribution at $\tau_e$ has a well defined maximum, the mass is not concentrated in a narrow interval around this maximum a posteriori (MAP) value. Given the fact that the model predictions obtained by simulating the model with parameters sampled from the posterior distribution are in excellent agreement with the experimental data and that the inference procedure previously yielded identifiable posterior distributions for the excitation timescale, $\tau_e$, the question of what causes the non-identifiability of the $\tau_e$ parameter, and what this implies for the applicability of the model, warrants a closer look. For this reason,

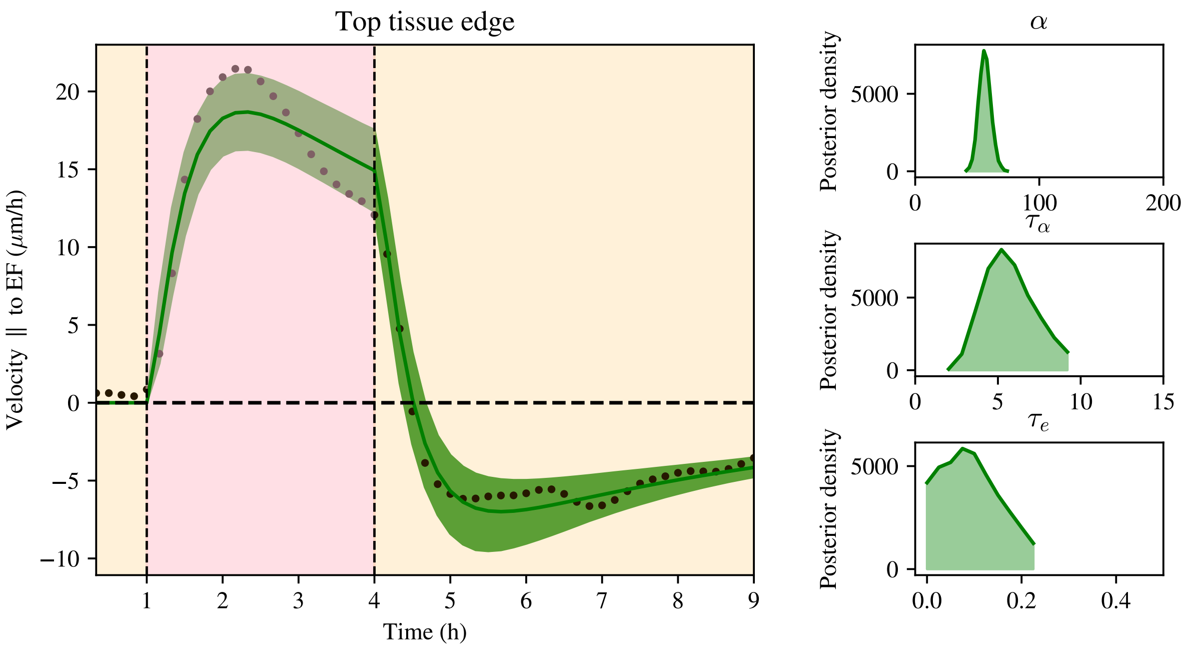


Figure B: Bayesian inference for the top edge velocity model in Equation (S3). Left panel: posterior 95% confidence interval for the top edge velocity model in green with experimental data (black scattered points). Right panel: posterior distributions for model parameters $\alpha, \tau_a$, and $\tau_e$, respectively, from top to bottom.

we turn to investigating the profile log-likelihood of the data under different model predictions with varying values of the model parameters.

To understand the influence of different values of the model parameters $\alpha$, $\tau_a$, and $\tau_e$ on the model predictions, we create a grid of size $50 \times 50 \times 50$ spanning the prior ranges shown for the distributions in Figure B. For each parameter value in the grid, we perform one forward simulation of the model in Equation (S3) and compute the sum of squared differences between the resulting model solution and the experimental data. Under the assumption of constant additive Gaussian noise, as was used in the MCMC implementations for Bayesian inference in this paper, the sum of squared differences is proportional to the log-likelihood. Therefore, by computing the sum of squared distances on the grid of values, we can interrogate the profile likelihoods for the different model parameters. These profile likelihoods, in turn, allow us to understand the effect of each of the three individual parameters on the model solutions. In Figure C we show the pairwise loss functions, which are proportional to the bivariate log-likelihoods, as well as the one-dimensional least squares profiles for each of the model parameters, which are equally proportional to the one-dimensional profile likelihoods.

The pairwise minimum loss landscapes in Figure C, which show the minimum sum of squared errors when marginalised over the third parameter, show that there is a well defined minimum

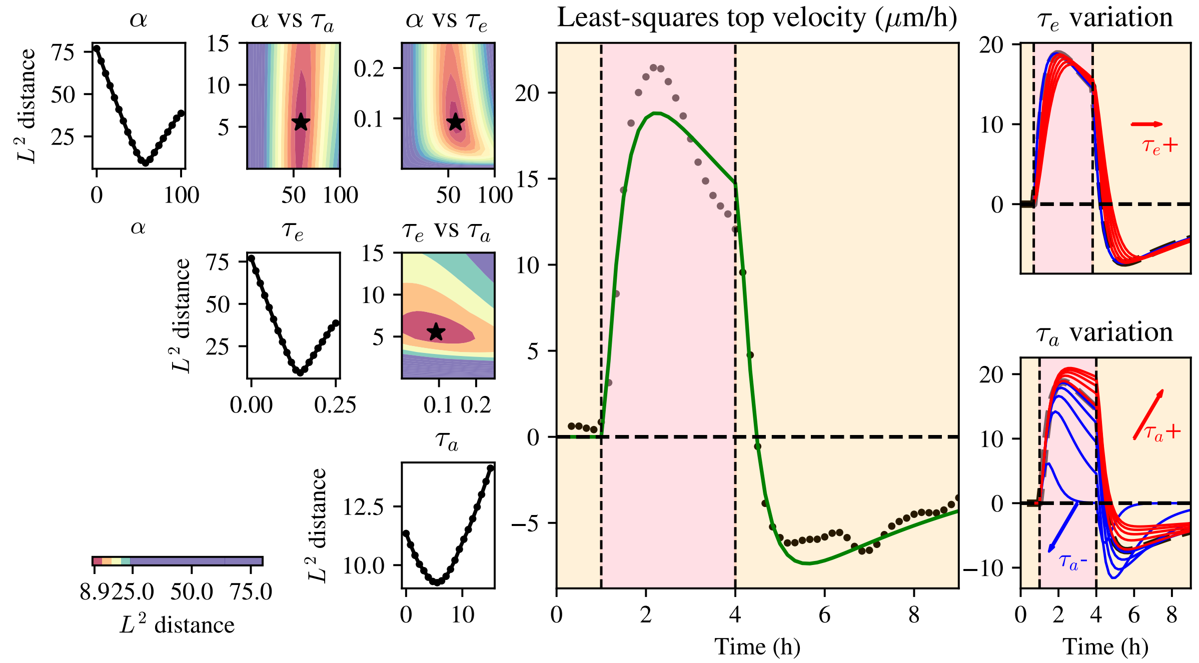


Figure C: Analysis of model solutions on a parameter grid. Left panel: pairwise loss landscapes shown off-diagonal, with the colourbar denoting the sum of squares between model solutions and parameter values (star denoting least squares parameter combination), marginal least squares profiles shown on diagonal. Middle panel: model solution corresponding to the least-squares parameter combination (green line) compared to experimental data (black points). Right panel: behaviour of model solutions as parameters $\tau_e$ and $\tau_a$ are varied.

value for the sum of squared errors. This is also made clear in the one-dimensional profiles in Figure C, which show the minimum sum of squared differences for each value of the different parameters. These profiles all have a clearly defined minimum, sum of squares error increasing rapidly upon deviations from the parameter value corresponding to least squares. Therefore, the profile likelihoods all point to a well-defined maximum likelihood, despite the posterior distributions in Figure B not being informative. For this reason, we ask why the posterior distribution fails to be informative of (some of) the model parameters, despite there being a well-defined maximum likelihood estimate of the parameters. The loss landscapes in Figure C, show that the model parameters $\tau_a$ and $\tau_e$ are *sloppy*. For a dynamical system, parameters are called sloppy if their values can be perturbed significantly without causing significant changes in the model output [4]. In the case of the squared distance profiles in Figure C, it can be seen that large deviations in $\tau_e$ from the minimum of the bivariate plots still lead to similar values of the sum of squared errors.

We illustrate how the model is sloppy by fixing the values of $\alpha$ , $\tau_a$, and $\tau_e$ at their maximum likelihood estimates from the the profiles in Figure C and simulating the model according to changing values of $\tau_a$ and $\tau_e$, as shown in the rightmost panel of Figure C. While variations in $\tau_a$ result in pronounced changes in both the shape and magnitude of the model solutions, variations in $\tau_e$ result in much smaller overall changes in the solution. In particular, reducing the value of $\tau_e$ leads to very small changes in the solution, with the rise of the velocity to its peak being nearly indistinguishable between different values of $\tau_e$ in the limit as $\tau_e \to 0$. We argue that, therefore, non-identifiability of the parameter $\tau_e$ in this case can be easily understood given the fact that these variations in model solutions are much smaller than the variance present in the experimental data. On a practical level, these issues arise because data acquisition in the experiment is only in 10 minute intervals;xt the data are not temporally resolved enough to tell different values of $\tau_e$ apart when there is only one time series. The inference results in Section 3 of the main text demonstrate that this issue does not occur in the presence of more data, but in the case of a single time series the temporal resolution becomes problematic.

In summary, we have shown that relaxing the assumption that the effective signal in the monolayer is always positive can explain the finding that the velocity in the top edge of the monolayer turns negative for several hours after the electric field is turned off. By carefully investigating model predictions with varying parameter values, we found that the undershoot is well-described by the maximum likelihood value of the parameters, which was difficult to identify in practice owing to the sloppiness in the model. Interestingly, such an undershoot is not observed any of the other tissue regions studied in this work, which begs the question for future work: why does signal transduction during electrotaxis appear to be intrinsically different in different tissue regions?

### References


[1] Abraham E Wolf, Matthew A Heinrich, Isaac B Breinyn, Tom J Zajdel, and Daniel J Cohen. Short-term bioelectric stimulation of collective cell migration in tissues reprograms long-term supracellular dynamics. *PNAS Nexus*, 1(1), 3 2022.

[2] Simon F Martina-Perez, Isaac B Breinyn, Daniel J Cohen, and Ruth E Baker. Optimal control of collective electrotaxis in epithelial monolayers. *arXiv preprint arXiv:2402.08700*, 2024.

[3] Jonathan Fouchard, Tom P J Wyatt, Amsha Proag, Ana Lisica, Nargess Khalilgharibi, Pierre Recho, Magali Suzanne, Alexandre Kabla, and Guillaume Charras. Curling of epithelial monolayers reveals coupling between active bending and tissue tension. *Proceedings of the National Academy of Sciences*, 117(17):9377–9383, 4 2020.

[4] Oana-Teodora Chis, Alejandro F Villaverde, Julio R Banga, and Eva Balsa-Canto. On the relationship between sloppiness and identifiability. *Mathematical Biosciences*, 282:147–161, 2016.
